# Structural insights into the binding mechanism of *Plasmodium falciparum* exported Hsp40-Hsp70 chaperone pair

**DOI:** 10.1101/311191

**Authors:** Ankita Behl, Prakash Chandra Mishra

## Abstract

Expression of heat shock proteins in *Plasmodium falciparum* (Pf) increases during febrile episodes to play key roles in several necessary cellular processes. ‘PFA0660w-PfHsp70-x’, an exported chaperone pair is known to co-localize to specialized intracellular structures termed J-dots, and has been implicated in trafficking of the major virulence factor, PfEMP1 (*Plasmodium falciparum* erythrocyte membrane protein 1) across the host cell. This article highlights for the first time detailed structural analysis of PFA0660w-PfHsp70-x chaperone pair to better understand their binding mechanism. Here, we have modeled reliable molecular structures for the complete conserved region of PFA0660w and PfHsp70-x. These structures were evaluated by different structure verification tools followed by molecular dynamics (MD) simulations. The model of PFA0660w was subjected to docking with PfHsp70-x using Haddock to reveal a number of residues crucial for their bipartite interaction, and also performed MD simulations on the complex. The peptide binding clefts of PFA0660w and its other *Plasmodium* species homologs were found to be bigger than their counterparts in higher eukaryotes like yeast, humans and *C. parvum*. Based on our results, we propose a model for PFA0660w-PfHsp70-x interaction and a mechanism of substrate binding, and compare it with its dimeric human counterparts. Owing to these striking structural differences between the host and parasite chaperones, such information on the essential Hsp40 and its partner Hsp70 may form the basis for rational drug design against fatal malaria.

## Introduction

Prophylaxis and chemotherapeutics, the potential ways for combating malaria disease has led to decrease in global mortality rate, yet developing resistance of malaria parasite to available drugs raises concerns about control of the disease [1]. Lack of a successful vaccine for malaria signifies the constant need for identification of novel drug targets for the development of anti-malarials. Molecular chaperones, specifically heat shock protein (Hsp) family form an important class of anti-malarial drug targets as their expression is upregulated in response to elevated body temperature of the human host. Therefore, they are considered key proteins that play crucial role for parasite survival during the febrile episodes [2]. Proteome analysis and loss of function mutant study revealed members of Hsp40 family to form a part of malaria exportome [3–7]. Presence of heat shock proteins in exportome underlines their importance in different aspects of malaria biology that includes protein folding and trafficking, translocation of exportome proteins across the parasitophorous vacuolar membrane and assembly and disassembly of multi-protein complexes [8–10]. The necessity of correctly folded essential proteins involved in host cell remodelling and sequestration further marks the importance of chaperones in parasite biology that accounts for about 2 % of the Pf genome [6].

Among various classes of molecular chaperones, Hsp40-Hsp70 interaction is considered the most ubiquitous chaperone protein partnership which facilitates folding of specific substrate protein in distinct subcellular localization [11]. Hsp40 function as cochaperones of Hsp70 and stimulate ATP hydrolytic rate which is necessary for driving their ATPase cycle [12]. The repeated cycles of interaction of Hsp70 with substrate peptide, facilitated by Hsp40 prevent substrate aggregation and ensure its proper folding [18]. Additionally, Hsp40s are suggested to govern the substrate specificity of Hsp70 and enable them to interact with diverse range of substrate proteins [19]. The Pf genome codes for about 49 Hsp40s and six Hsp70s [20, 21]. Since the number of Hsp40 proteins far exceeds that of Hsp70s, it is suggested that Hsp70 proteins are regulated by multiple Hsp40 partners [12]. Hsp40 members are characterized by the presence of J domain containing the His-Pro-Asp (HPD) tripeptide that forms the signature motif of this protein family [12]. Out of 49 PfHsp40 proteins, 36 members have a signature J-domain and are characterized into three different classes (Type I-III). Additionally, 13 Hsp40 proteins are with a J-like domain where the HPD motif is modified, and are included in type IV class [20]. Type I Hsp40s have typical characteristics of prokaryotic DnaJ and carry N-terminal J-domain, flexible ‘G/F’ region (50-100 amino acid), a cysteine-rich region (consisting of four CXXCXGXG repeats forming a zinc-finger domain) followed by a less conserved C-terminal domain (120–170 amino acids). Type II Hsp40s are similar to type I but lack the cysteine rich region; however they are believed to be functionally equivalent to type I. Type III Hsp40s exhibit only the signature J domain, not necessarily to be present at N terminal. Three PfHsp40s (PFA0660w, PF11s_0034 and PF11_0509) contain PEXEL motif required for protein export to erythrocyte cytosol, and are supposed to be essential as per gene knock out studies [7]. Hsp70s possess two domains that include a 45 kDa N-terminal nucleotide-binding domain (NBD) having ATPase activity, and a 25 kDa substrate binding domain joined by a linker region [17, 22]. Biochemical and structural data suggests that N terminal NBD of Hsp70 forms the major binding site for highly conserved J domain of Hsp40 [23].

Experimental evidences suggest the existence of Hsp40-Hsp70 interactions in malaria parasite that play key role in protein folding mechanisms. ‘PfHsp70-1’, a nucleo-cytoplasmic resident protein is regulated by several Hsp40s. A type I Hsp40 (PF14_0359) localizes in the parasite cytosol and functionally interacts with PfHsp70-1, [24]. Pesce *et al*. revealed Pfj4 (PFL0565w, type II Hsp40) to be present in common complex with PfHsp70-1 [25]. A more recent data providing the evidence for functional chaperone-cochaperone partnership within the parasite cytosol was shown between PfHsp70-1 and ‘PFB0595w’, a type II Hsp40 protein [26]. Hsp40 proteins have also been reported to play important role in providing pathogenicity to *P. falciparum*. Formation of knobs that mediate the process of cytoadherence is considered pivotal to pathophysiology of the disease. KAHRP, MESA/PfEMP-2, PfEMP-1 are reported to be involved in knob formation in infected erythrocytes [27]. A study reports that an exported type II Hsp40 (PF3D7_0201800) co-localise with both KAHRP and PfEMP1, suggesting its possible association in knob mediated pathogenesis [28].

PFA0660w has been reported to function as co-chaperone of PfHsp70-x and these co-localize to mobile intracellular structures called J-dots in the cytosol of infected RBCs [29, 30]. Protein binding studies by Daniyan et al. revealed *in vitro* interaction of PFA0660w with PfHsp70-x [29]. The present study sought to propose a three dimensional structure of complete conserved region of PFA0660w to understand its substrate binding characteristics and interactions with PfHsp70-x. Using modeled structures of PFA0660w and PfHsp70-x, docking studies were conducted to explore crucial residues involved in their interaction. The proposed PFA0660w-PfHsp70-x docked complex was evaluated using molecular dynamics simulation followed by post MD analysis such as RIN and Dimplot. These studies provide detailed structural information in context of PFA0660w-PfHsp70-x interaction which may help in targeting these proteins and their docked complex towards future anti-malarial drug design.

## Material and methods

### Structure prediction and validation

Sequence of all proteins used in the study were retrieved from PlasmoDB database [31]. Multiple sequence alignment was performed using Clustal omega [32] whereas Blastp was used to find the homologs of PFA0660w [33]. For moleculer modeling, Blastp against the RCSB Protein Databank was used to find the suitable template [34]. Different approaches were made to get a good quality model of complete conserved region of PFA0660w (amino acid sequence range: 81 to 392) using Swiss model tool and Modeller9.14 [35,36]. Basic modeling in Modeller 9.14 was performed using the structure of *Thermus thermophilus* DnaJ as a template (PDB ID: 4J80) whereas in advanced modeling, multiple templates (PDB ID’s 4J80, 3LZ8, 2Q2G and 1BQZ) were used for model building. DaliLite v.3 was used for multiple structural alignments [37]. C-terminal region of PFA0660w and its homologs in Pf3D7 and in other strains of *Plasmodium* including *P. vivax, P. yoelii and P. berghei* share high level of sequence similarity with dimerization domain of *C. parvum* Hsp40 ‘Cgd2_1800’. The atomic resolution structure of Cgd2_1800 (PDB ID: 2Q2G) was used as a template to build their model structures by Swiss model [35]. For PfHsp70-x, crystal structure of bovine Hsc70 having 75% sequence identity was used for generating the model structure using Modeller9.14 [36]. All predicted models were refined using 3Drefine server for improving the qualities of protein structures [38] and were evaluated using Verify 3D, Errat and Rampage programs [39–41].

### Protein-protein docking and identification of interface residues

The Easy interface of Haddock webserver was used for molecular docking studies [42]. No passive residues were set for docking. The best docked structure was selected on the basis of Haddock score, the weighted sum of the Van der Waals energy, electrostatic energy, desolvation energy, the energy from restraint violations and the buried surface area. Docked complexes were evaluated for their total stabilizing energy and normalized energy per residue using PPCheck webserver [43]. The Protein Interactions Calculator (PIC) webserver was used to identify the possible interface residues involved in interactions in the docked structure [44]. While submitting the structures to PIC server, default settings were used.

### Molecular dynamics simulations

MD simulations for PFA0660w and PfHsp70-x models and their docked complex were performed using the program NAMD (Nanoscale Molecular Dynamics program; v 2.12) [45] and visual molecular dynamics (VMD 1.9.3) [46]. The protein complex was solvated with a water box using CHARMM (Chemistry at HARvard Macromolecular Mechanics) 22 parameter file [47]. Electrostatic interactions were calculated by the Particle Mesh Ewald method with grid dimensions of 1.0 Å [48]. The system was subjected to energy minimization and production run of 2 ns was carried out by applying periodic boundary conditions, under constant pressure (1 atm) and temperature (310 K). The pressure and temperature were controlled by the Langevin dynamics [49]. The trajectory files generated by the MD simulation of PFA0660w-PfHsp70-x complex were analysed for root mean-square deviation (RMSD), root-mean square fluctuation (RMSF) and radius of gyration (Rg) using VMD Tk-Console command. The number of inter-chain hydrogen bonds (NH-bond) formed by PFA0660w-PfHsp70-x complex during the simulation was calculated using VMD HBonds Plugin. The NH-bonds were determined based on the cutoffs for the angle (30°) and distance (0.35 nm) between donor and acceptor heavy atoms.

### Protein-protein interaction analysis

A representative structure was selected from the trajectories obtained after MD simulation using cluster analysis. Clustering of trajectories was performed based on pairwise best-fit root-mean-square deviations (RMSDs) using MD Movie, a tool in the MD/Ensemble Analysis category in UCSF Chimera [50]. Average structure of the most populated cluster was selected for post MD protein-protein interaction analysis. Residue Interaction Network (RIN) Analysis is a technique that represents various interactions in the form of a detailed network model. The RIN profile of obtained representative structure of PFA0660w-PfHsp70-x complex was generated using RING 2.0 web server [51] and analysed using Cytoscape 3.5.1 [52]. The nodes in a RIN profile represent backbone amino acids whereas, the edges correspond to distinct inter residue interactions. Profiles of interacting amino acid residue pairs of representative structure of PFA0660w-PfHsp70-x complex was further analysed using the dimplot command of LigPlot+ (v 1.4.5) [53]. DIMPLOT program generates schematic 2-D graphical representations of residue-residue interactions across the interface of protein-protein complexes.

### Peptide binding cleft prediction and analysis

Peptide binding cleft parameters of PFA0660w, its homologs in *Plasmodium* and crystal structure of type II Hsp40 of human (PDB ID: 2QLD), yeast (PDB ID: 1C2G) and *C. parvum* (2Q2G) were computed using DoGSiteScorer: Active Site Prediction and Analysis Server [54]. Among various cavities predicted for the structures, peptide binding cleft responsible for capturing the substrate peptide was identified by considering the corresponding residues of yeast Sis1 (V184, L186, F201, I203 and F251) reported to be involved in forming the substrate binding cleft [55].

## Results

### Homology modeling, MD simulation and evaluation

The main objective of this study is to understand the binding mechanism of PFA0660w-PfHsp70-x chaperone pair. For this, we modeled three dimensional structures for the complete conserved region of PFA0660w and PfHsp70-x using comparative modeling and employed protein–protein docking techniques. Based on the complex obtained, we explored the interaction interface through molecular dynamics simulations and identified hotspot residues using RIN and Dimplot.

Blastp against PDB database using PFA0660w sequence showed significant matches for J domain and C-terminal region whereas only one match of *Thermus thermophilus* DnaJ (PDB ID:4J80; 27% identity) was identified for the complete conserved region of protein (81-392 amino acids) (Table 1). The conserved region includes J domain followed by G/F region and C terminal substrate binding domain. Using 4J80 as template, we modeled the three dimensional structure of PFA0660w by Swiss model and Modeller9.14. Various tests like Verify 3D, Errat and Ramachandran plot were run on the generated models to assess their quality and were found not suitable owing to their low scores (Table 2). Verify 3D analysis calculates the compatibility of an atomic model (3D) with its own amino acid sequence (1D). A model is considered to be of good quality if more than 80% residues have average 3D-1D score>=0.2. ERRAT analyses the statistics of non-bonded interactions between different atom types, and depicts the overall model quality. Higher scores indicate higher quality, however generally accepted range for the good quality model is > 50. Even after refining the structures with 3Drefine, the models were observed to have low quality (Table 2). 3Drefine follows a two-step refinement protocol based on optimizing hydrogen bonding network along with atomic-level energy minimization for improving the local and global structural quality measures [38].

**Table 1:** Templates used for modeling PFA0660w with their PDB IDs, percentage of query cover and sequence similarity.

| Source organism | PDB ID | % Query cover | % sequence identity |
| --- | --- | --- | --- |
| Crystal structure of dimerization domain of Hsp40 from *C. Parvum* (Cgd2_1800) | 2Q2G | 43 | 49 |
| EPR-X-RAY diffraction structure of *Thermus Thermophilus* DnaJ | 4J80 | 78 | 27 |
| Structure of putative chaperone DnaJ from *Klebsiella Pneumoniae* | 3LZ8 | 60 | 24 |
| NMR structure of *E.coli* DnaJ domain | 1BQZ | 19 | 53 |

**Table 2:**
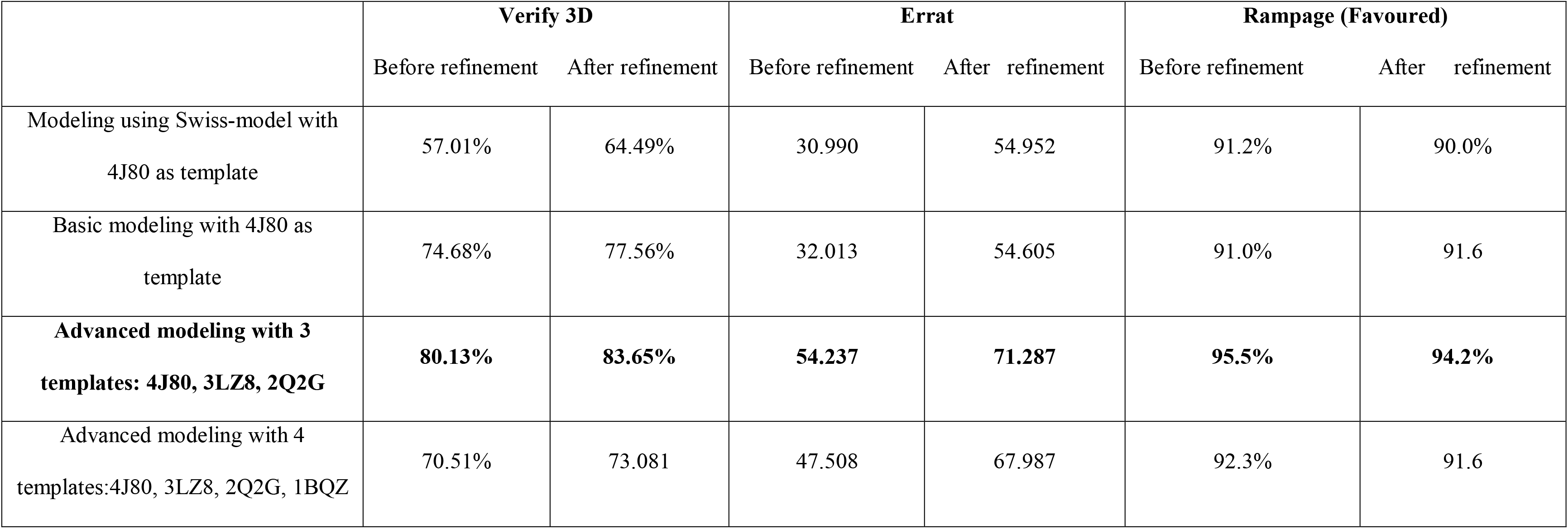
Verify 3D, Errat and Ramachandran scores for the validation of predicted models using different approaches. Validation scores in bold represent the best model selected for PFA0660w.

|  | **Verify 3D** |  | **Errat** |  | **Rampage (Favoured)** |  |
| --- | --- | --- | --- | --- | --- | --- |
|  | Before refinement | After refinement | Before refinement | After refinement | Before refinement | After refinement |
| Modeling using Swiss-model with 4J80 as template | 57.01% | 64.49% | 30.990 | 54.952 | 91.2% | 90.0% |
| Basic modeling with 4J80 as template | 74.68% | 77.56% | 32.013 | 54.605 | 91.0% | 91.6 |
| **Advanced modeling with 3 templates: 4J80, 3LZ8, 2Q2G** | **80.13%** | **83.65%** | **54.237** | **71.287** | **95.5%** | **94.2%** |
| Advanced modeling with 4 templates:4J80, 3LZ8, 2Q2G, 1BQZ | 70.51% | 73.081 | 47.508 | 67.987 | 92.3% | 91.6 |

Therefore to build a reliable model for the complete conserved region of PFA0660w (81-392 amino acids), we employed multiple template based approach. Four templates from the Blastp results were selected based on query coverage and sequence identity (Table 1). The model structure generated using three templates (4J80, 3LZ8, 2Q2G) was refined and evaluated by different structure verification tools and was found to be sufficiently robust for structural analysis. In the Verify 3D analysis, none of the amino acids was observed to have a negative score (Fig.1a). 84.97 % of the residues had an average 3D-1D score of <0.2, depicting that the predicted model was compatible with its amino acid sequence. The amino acid environment was evaluated using ERRAT plot which showed an overall quality factor of 71.287 (Fig.1a). The stereo-chemical evaluation of backbone psi and phi dihedral angles was carried out with Ramachandran plot analysis using Rampage. 94.2% of the residues were observed to fall within the favoured region, indicating the accuracy of backbone dihedral angles (Fig.1a). It is generally accepted that a score close to 100% depicts good stereochemical quality of the model. Therefore, these Rampage results suggesting 94.2% score indicate that the predicted model is of good quality. Structure based sequence alignment using Dali server shows PFA0660w model to be similar to templates structures used in modeling. We also attempted to get a more reliable model by adding another template (1BQZ) for the J domain in advanced modeling. However, addition of this template led to decrease in quality of resulting structure; therefore it was excluded for modeling PFA0660w. The comparison of Verify 3D, Errat and Ramachandran scores for predicted models using different approaches is given in Table 2.

**Fig. 1.**
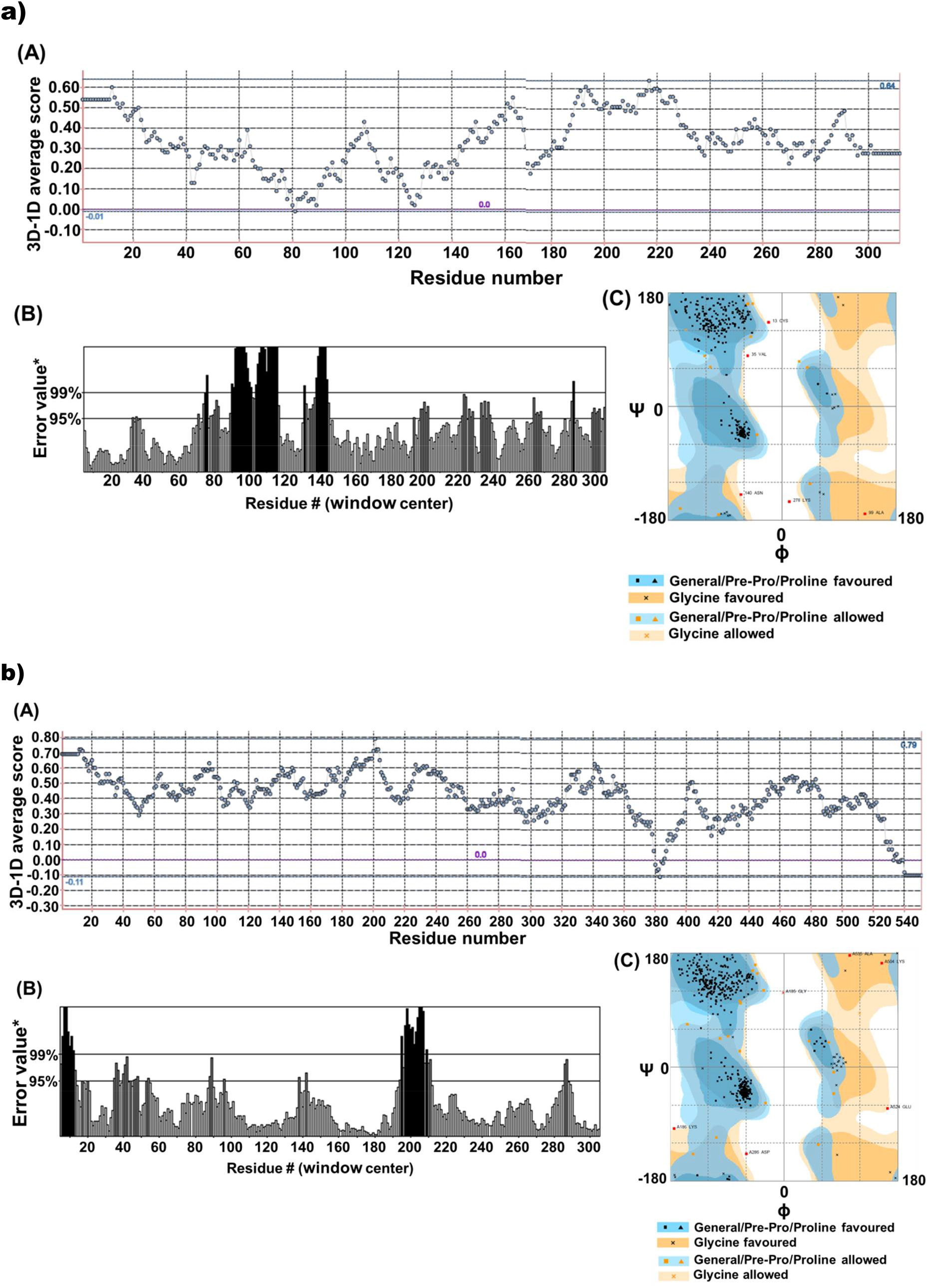
Verify 3D (A), Errat (B) and Ramachandran plots (C) of modelled (**a**) PFA0660w and (**b**) PfHsp70-x.

We modeled the complete conserved region of PfHsp70-x (35 to 585 amino acid residues) using crystal structure of bovine Hsc70 (75% sequence identity) as template. The predicted model was refined and evaluated using Verify 3D, Errat and Rampage programs. The Ramachandran plot of the refined model showed 95.4% of the residues to lie in the most favoured region (Fig.1b). Verify 3D and Errat score were 92.20% and 86.137 respectively (Fig.1b). These scores fit well within the range of a high quality model; therefore it was selected for further docking studies.

Molecular dynamics simulations were performed on selected models for individual proteins (PFA0660w, PfHsp70-x) to explore their internal dynamics. RMSDs of the protein backbone as compared with the initial conformation plotted with respect to simulation time suggested the structural models to be stable (Fig. 2a, upper panel). We also computed radius of gyration (Rg) as an indicator of protein structure compactness. A relatively steady Rg value indicates that proteins are stably folded (Fig. 2a, lower panel).

**Fig. 2:**
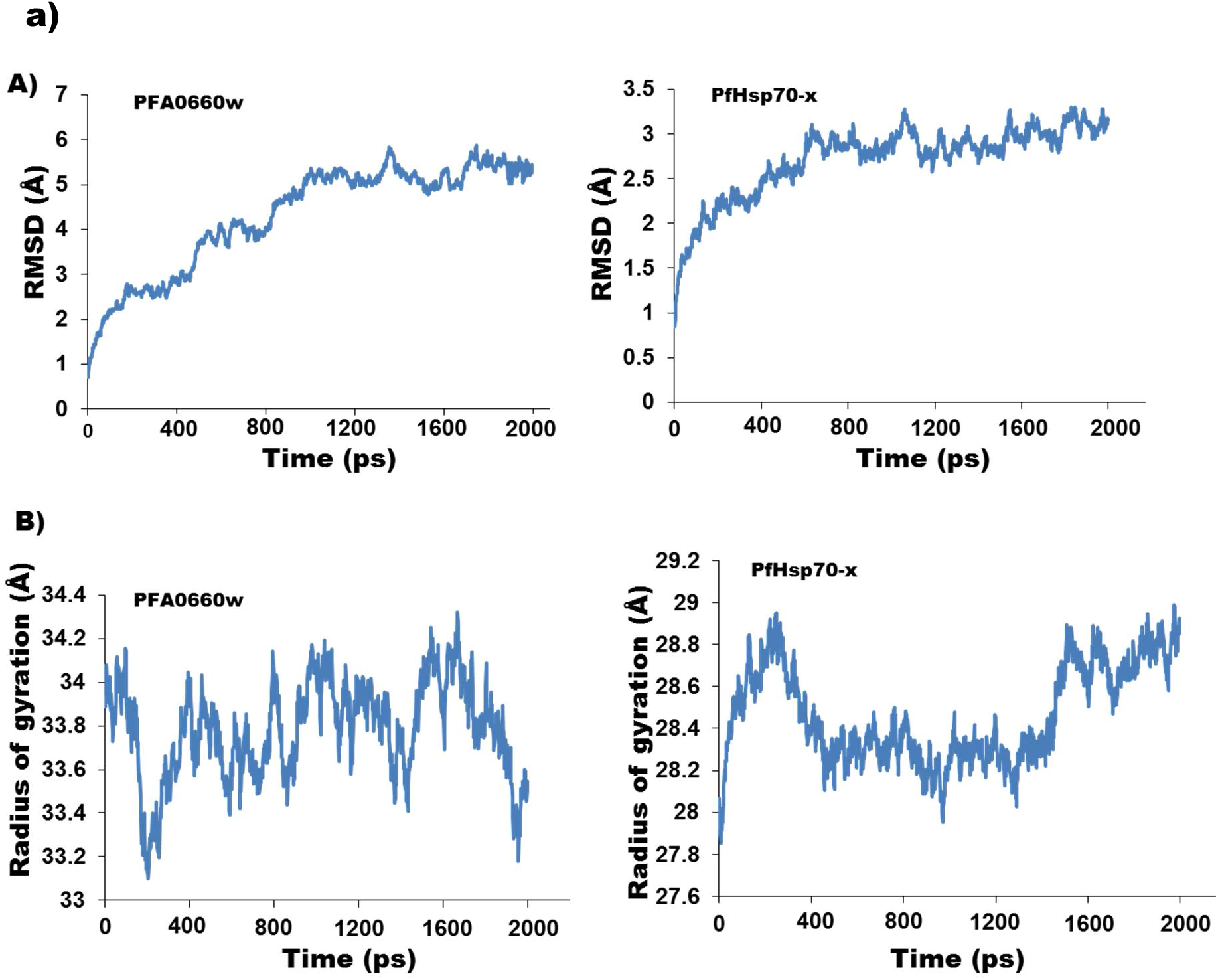
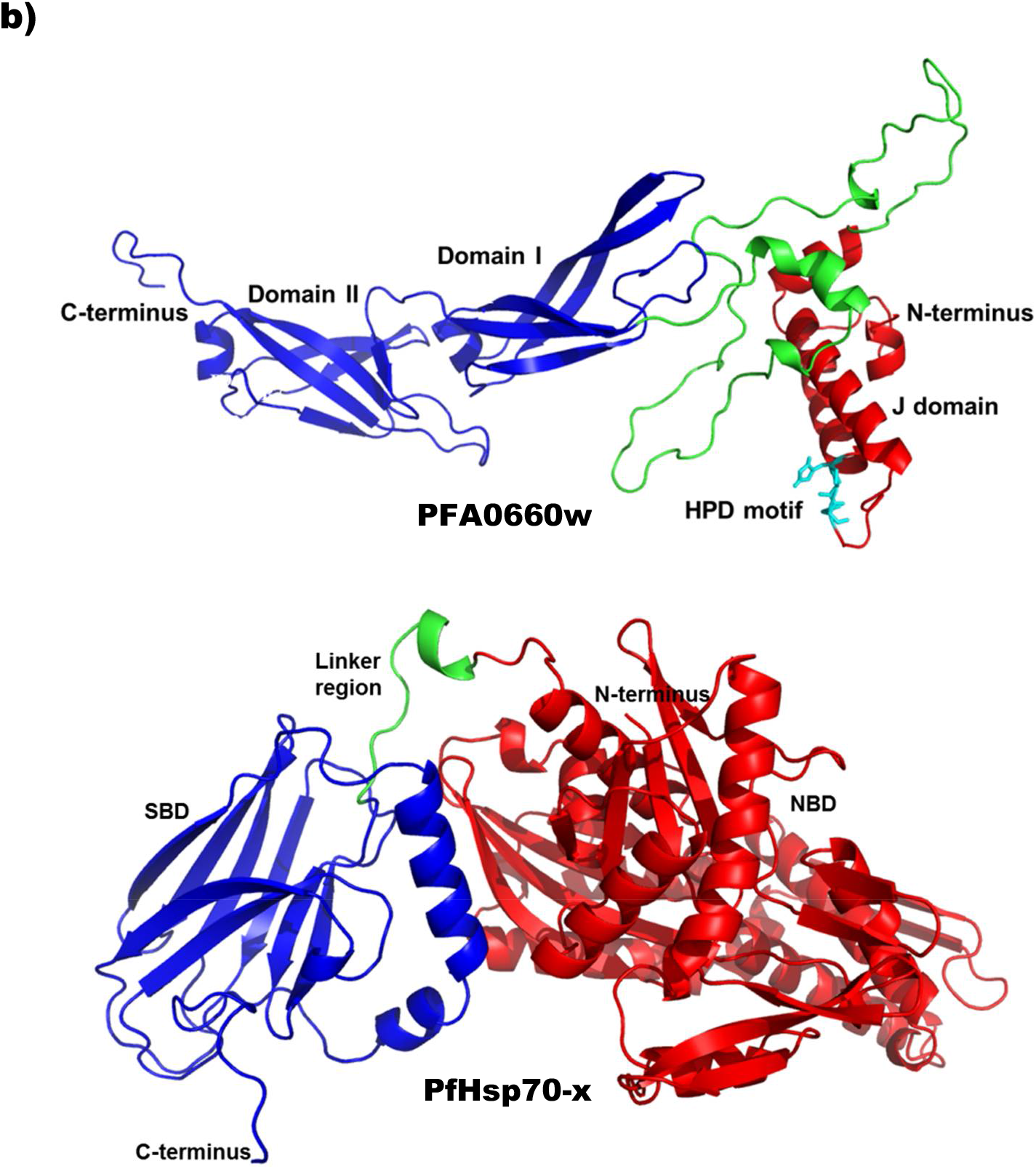
**a**) RMSD and radius of gyration graphs for PFA0660w and PfHsp70-x. (A) Plots of RMSD and (B) radius of gyration (Rg) for PFA0660w (left panel) and PfHsp70-x (right panel) for 2 ns MD simulations. **b**) Cartoon representation of homology model of PFA0660w (upper panel) and PfHsp70-x (lower panel). J domain (red) of PFA0660w comprises four alpha helices with HPD motif (shown in sticks and colored cyan), residing in the loop region between helix II and helix III. G/F region (green) mainly consists of loops. C-terminal peptide binding fragment (blue) carry two domains consisting of beta strands. Hydrophobic substrate binding cavity resides in domain I. PfHsp70-x carries N-terminus NBD (red) which is connected to SBD (blue) via linker region (green).

Structural analysis showed that J domain of PFA0660w comprises four α helices with HPD motif residing in the loop region between helix II and III (Fig. 2b, upper panel). G/F rich region consists mainly of loops with short α helices in between whereas C terminal region has two distinct domains consisting of eleven beta strands and two short α helices. Domain I forms the substrate binding site (Fig. 2b, upper panel). PfHsp70-x carries N-terminus NBD which is connected to C-terminus SBD via linker region. NBD comprises both α helices and beta strands while SBD mostly carry beta strands (Fig. 2b, lower panel).

### Molecular docking

Previous reports suggest that conserved HPD tripeptide motif of J domain mediate DnaJ interaction with ATPase domain of DnaK [18, 23]. Residues of DnaK reported to be critical in binding include R167, N170 and T173 [56]. Structure of auxilin J domain complexed with bovine Hsc70 revealed that I216 and V388 form a part of interface in Hsp40-Hsp70 complex [57, 58]. A more recent report deciphering the mechanisms of DnaJ-DnaK interactions revealed that it is the positively charged residues on helix II of DnaJ that form electrostatic interactions with the ATPase domain, particularly with negatively charged residues on segment 206-221 of DnaK [59].

Modeled structure of PFA0660w was docked with PfHsp70-x using Haddock to investigate their interaction interface. Protein–protein docking was performed by considering the corresponding residues entailed in previous reports as active sites [18, 23, 56–58]. Detailed account of these residues is represented in table 3. We selected docked PFA660w-PfHsp70-x complexes on the basis of their Haddock score. These complexes were evaluated for their total stabilizing energy and normalized energy per residue using PPCheck server (Table 4). The most stable docked complex (named as 1 Dock) with −263.83 kJ/mol of stabilizing energy was selected for structural studies. Normalized energy per residue for this complex was −3.26 kJ/mol. A standard energy range of −2kJ/mol to −6kJ/mol signifies correct docking pose by PPCheck. A number of potential interactions including hydrophobic, ionic interactions, and hydrogen bonds were computed for the selected docked complex (1 Dock) by PIC webserver. Detailed analysis revealed that interaction of PFA0660w with PfHsp70-x is bipartite in nature. J domain of PFA0660w binds with NBD and SBD of PfHsp70-x while residues of G/F region bind to SBD only (Fig. 3A).

**Fig. 3.**
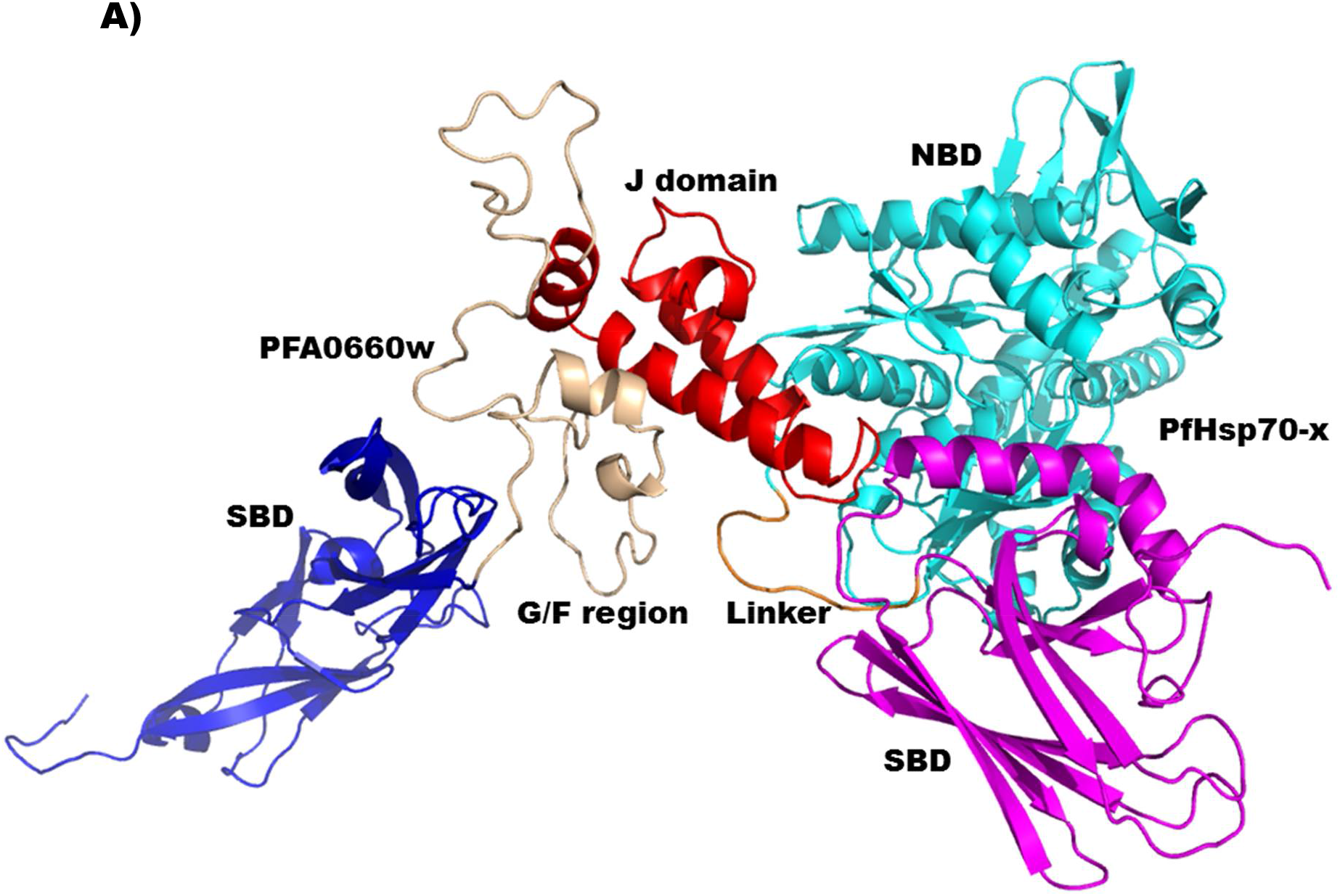
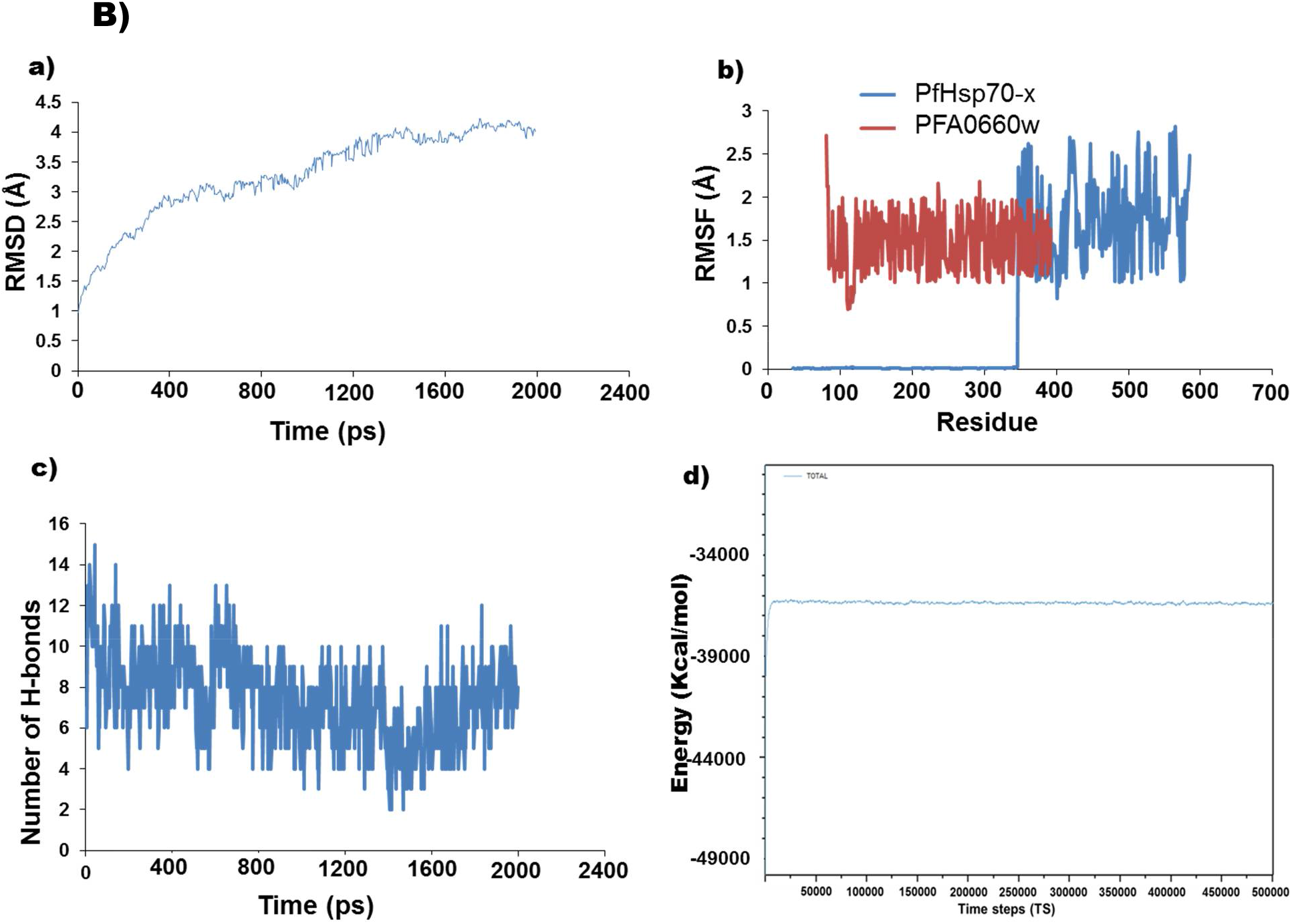
(**A**) Docked complex showing the interaction interface for PFA0660w and PfHsp70-x. J domain (red) of PFA0660w binds with NBD (cyan) and SBD (magenta) of PfHsp70-x whereas its G/F region (wheat) interacts solely with SBD (magenta) of PfHsp70-x. SBD of PFA0660w and linker region of PfHsp70-x are shown in blue and orange respectively. (**B**) Dynamic changes of PFA0660w-PfHsp70-x complex structure. (**a**), (**b**) RMSD and RMSF plots for the backbone of protein-protein complex respectively. (**c**) Inter-chain H-bond pattern during the period of simulation. (**d**) NAMD plot for total energy over time steps for the equilibrated system.

**Table 3:** Active site residues of PFA0660w and PfHsp70-x given to guide in docking. H110, P111 and D112 form the signature tripeptide motif whereas R99, R100, K104, K108 constitute positively charged residues in helix II of J domain of PFA0660w. E243, D244, E248 are negatively charged residues found in stretch of 207-218 amino acids of PfHsp70-x that corresponds to 206-211 of ATPase domain of DnaK. R201, N204, T207 of PfHsp70-x are the corresponding residues for R167, N170 and T173 of DnaK whereas I246 and V419 are the corresponding residues for I216 and V388 of bovine Hsc70 reported to be crucial for binding with Hsp40.

| Docked complex | Active site residues of PFA0660w | Active site residues of PfHsp70-x |
| --- | --- | --- |
| Dock 1 | R99, R100, K104, K108, H110, P111, D112 | R201, N204, T207, E243, D244, I246, E248, V419 |
| Dock 2 | R99, R100, K104, K108 | E243, D244, E248 |
| Dock 3 | H110, P111, D112 | R201, N204, T207, E243, D244, I246, E248, V419 |
| Dock 4 | H110, P111, D112 | R201, N204, T207, I246, V419 |

**Table 4:** Evaluation of docked complexes on the basis of their total stabilizing energy and normalized energy per residue. Docked complex shown in bold was selected for structural studies.

| Docked complex | Total stabilizing energy (kJ/mol) | Normalized energy per residue (kJ/mol) |
| --- | --- | --- |
| **Dock 1** | **-263.86** | **-3.26** |
| Dock 2 | -176.55 | -2.05 |
| Dock 3 | -215.70 | -2.88 |
| Dock 4 | -232.48 | -3.02 |

To infer conformational fluctuations, MD simulation for PFA0660w-PfHsp70-x complex was performed for 2ns using NAMD 2.12 package. The stability of the protein-protein complex relative to its initial conformation was determined by RMSD deviations of C-alpha backbone produced during the course of its simulation. The MD simulation for system shows a gradual increase in the RMSD value with fluctuations, stabilizing at an average of 4 Å (Fig. 3B, upper left panel). Further, the RMSF values of carbon alpha for each amino acid residue were obtained from the trajectory data of complex (Fig. 3B, upper right panel). We also computed inter-chain hydrogen bonds for the complex which was observed to be relatively stable for the entire period of simulation (Fig. 3B, lower left panel). NAMD plot for the total energy was plotted over time steps which reveal that total energy converges to an average value for the equilibrated system (Fig. 3B, lower right panel). A total of 6 structural representative snapshots were obtained at time intervals of 0.4 ns and the whole trajectory is represented in Fig. 4. These snapshots and trajectory movie of 2ns simulation (Movie S1) clearly demonstrate that PFA0660w interact with PfHsp70-x with two distinct binding sites viz. J domain and G/F region.

**Fig. 4.**
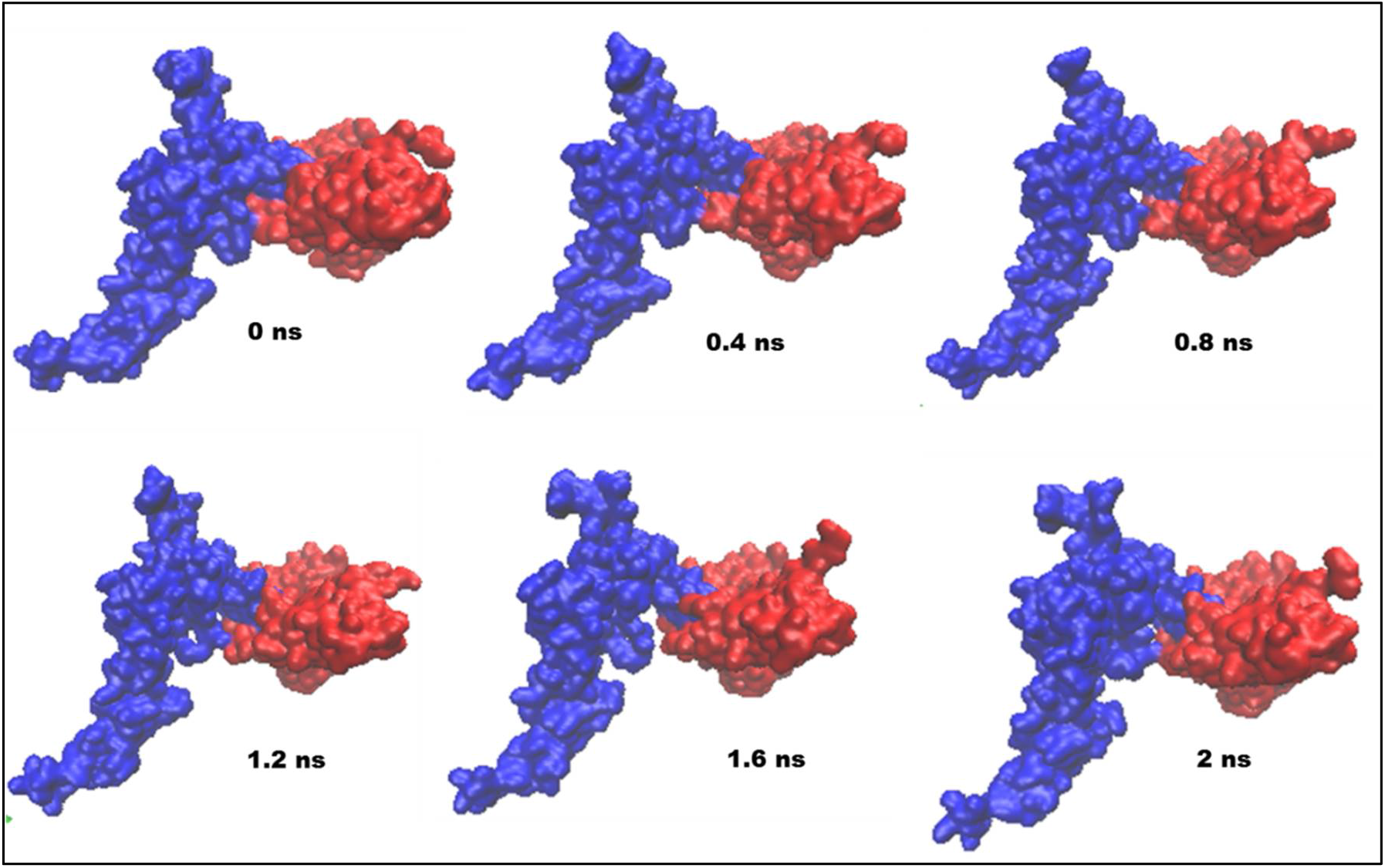
Snapshots of PFA0660w-PfHsp70-x docked complex from 2ns MD simulation. Blue and red correspond to PFA0660w and PfHsp70-x respectively.

### PFA0660w-PfHsp70-x complex interface analysis

The residue interaction network (RIN) analysis is used to identify key residue interactions in protein structures. It provides a visual interface to analyse the stability of connections formed by amino acid residues at the contact sites [51]. Clustering of trajectories was performed for selecting a representative structure for post MD analysis. Average structure of the most populated cluster was selected to identify hotspot residues using RING 2.0 webserver and Dimplot. Detail analysis of RIN plot and Dimplot reveals that residues of J domain of PFA0660w form stable interactions with NBD and SBD of PfHsp70-x. This includes Leu103, Met107, Lys108, His110, Asp112, Lys113, His114, Val115, Asn 116, Lys117, Gly118, Lys120 and Val121. Interface residues of PfHsp70-x involved in interaction network include Asp216, Lys 217, Asp244, Phe247, Gln407, Ile410, Leu411, Asp414, Gln415, Ser416 Ala418, Val419, Asp421, Leu422, Leu425, Gly539, Ser542, Lys 543 Asp544 and Asp545. His114 of loop region between helix II and III of J domain form maximum number of interactions. His114 forms hydrogen bonds with Asp216 (NBD) and Van der Waals interactions with Lys217 (NBD), Ser416 (SBD), Ala418 (SBD) and Lys543 (SBD) of PfHsp70-x. Lys113 and Lys108 of J domain also stabilize the interaction network by forming hydrogen bonds with Asp244 and Asp216 respectively of NBD of PfHsp70-x. Apart from residues of J domain, a residue of G/F region (Asn197) of PFA0660w was observed to form stable network with residues of SBD (Asp421, Leu422) of PfHsp70-x. Asn197 form hydrogen bonds, hydrophobic and Van der Waals interactions with Asp421 of PfHsp70-x. Asn197 also forms Van der Waals interactions with Leu422 and hydrophobic interactions with Lys420 of PfHsp70-x. Pictorial representation of all the residues involved in interaction interface between PFA0660w-PfHsp70-x complex is shown in RIN plot and Dimplot (Fig. 5A, B). Overall, analysis suggest that although, PFA0660w interacts primarily with PfHsp70-x through J domain, G/F region forms another binding determinant of PFA0660w-PfHsp70-x interaction.

**Fig. 5.**
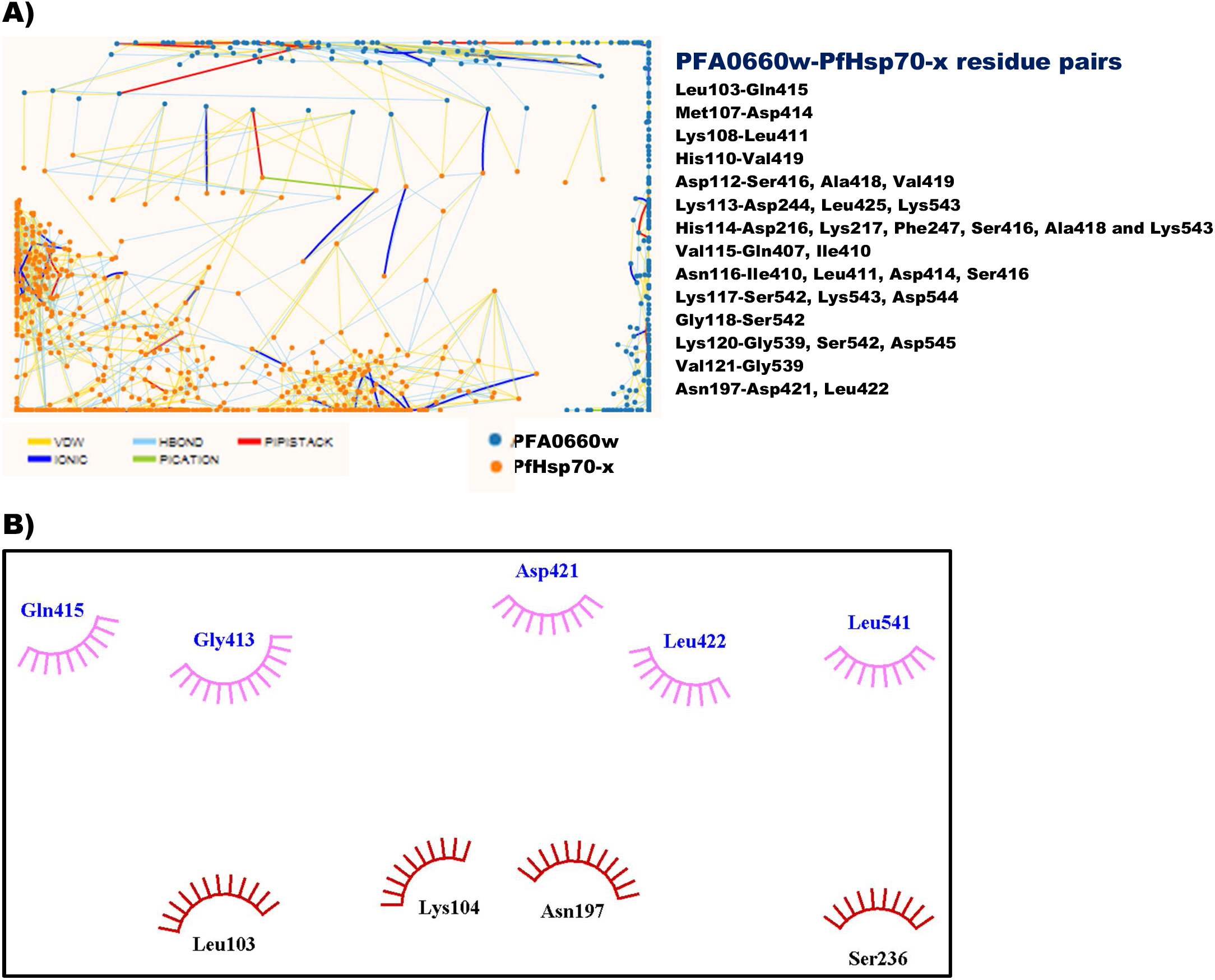
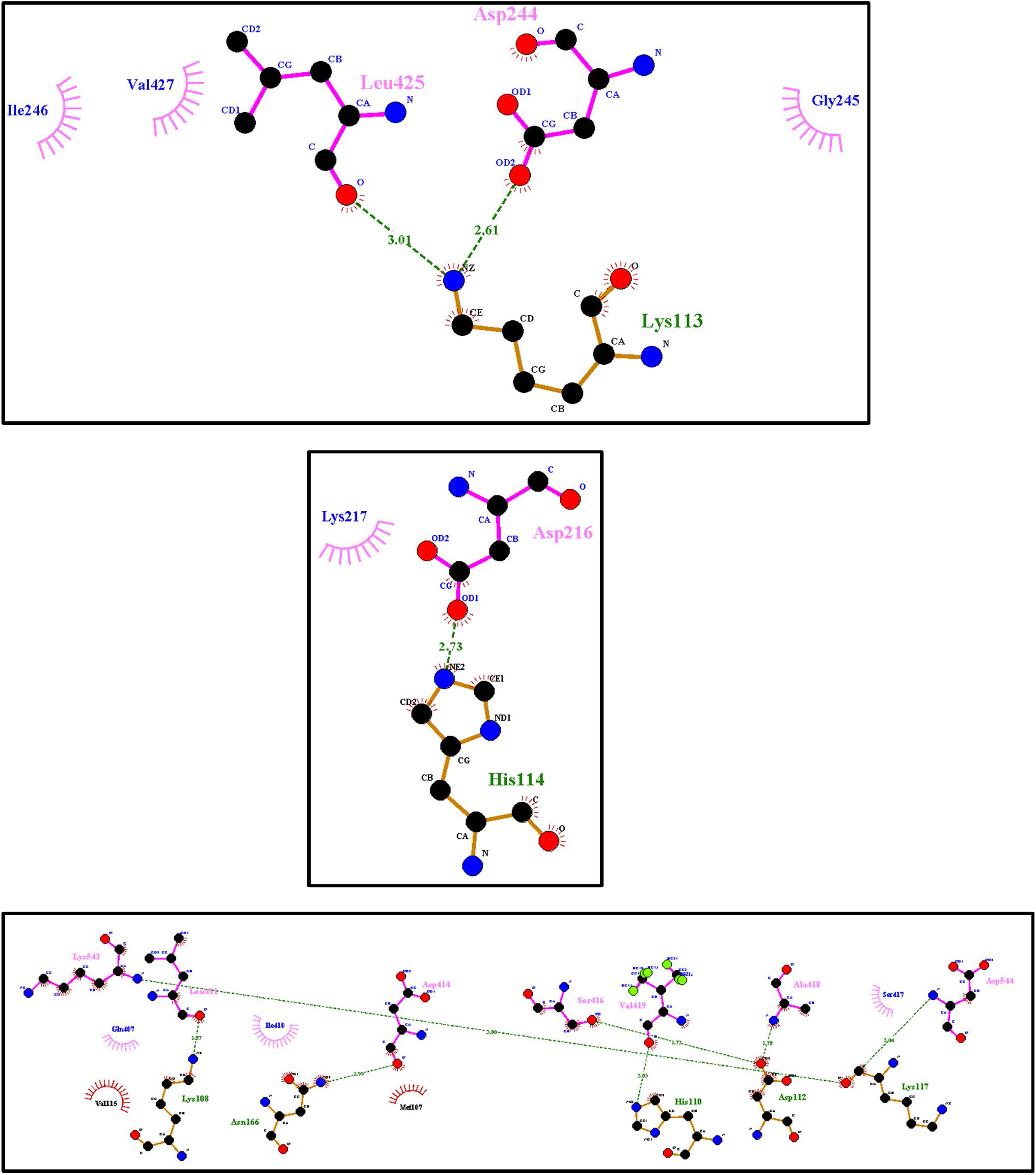
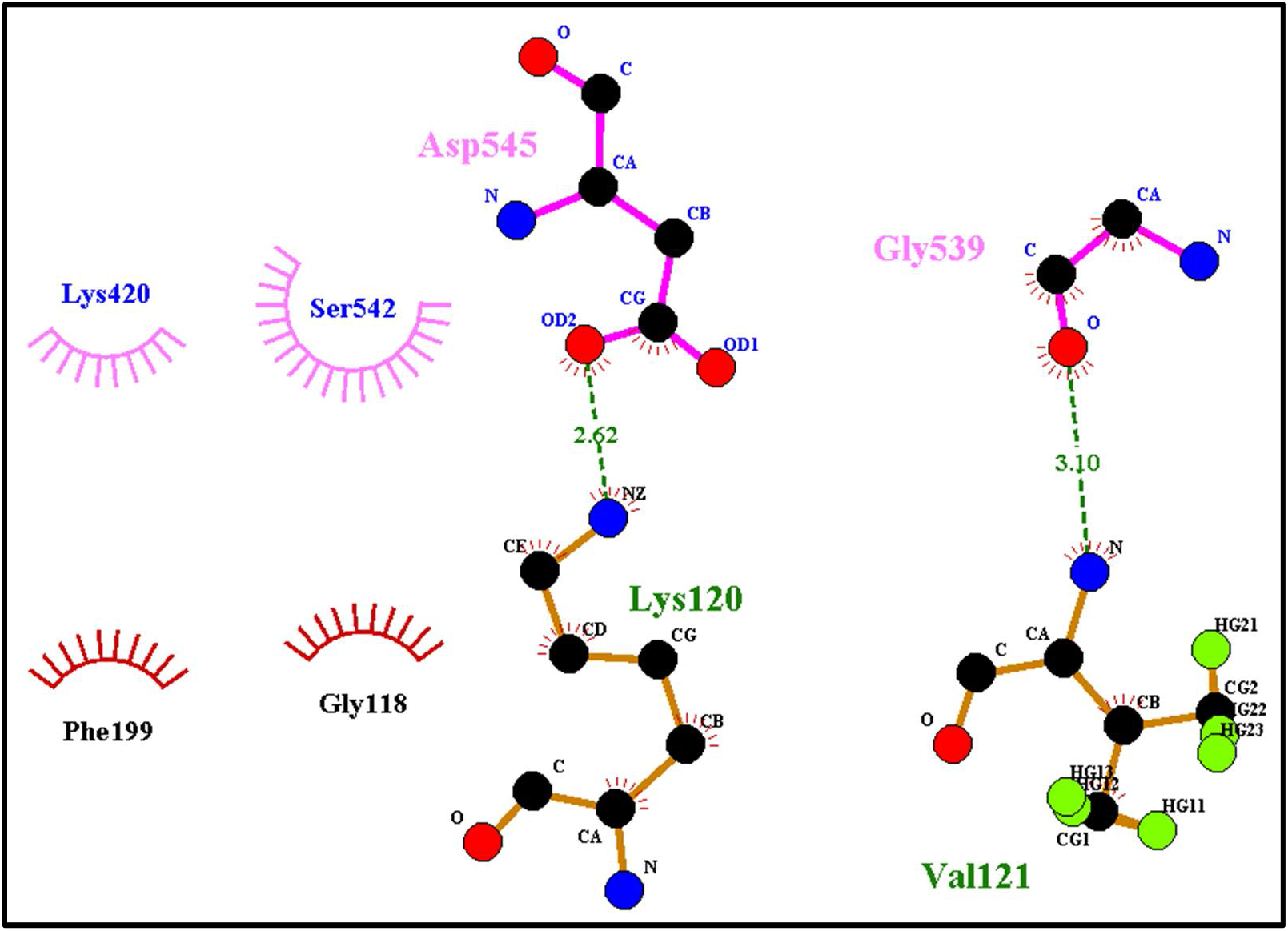
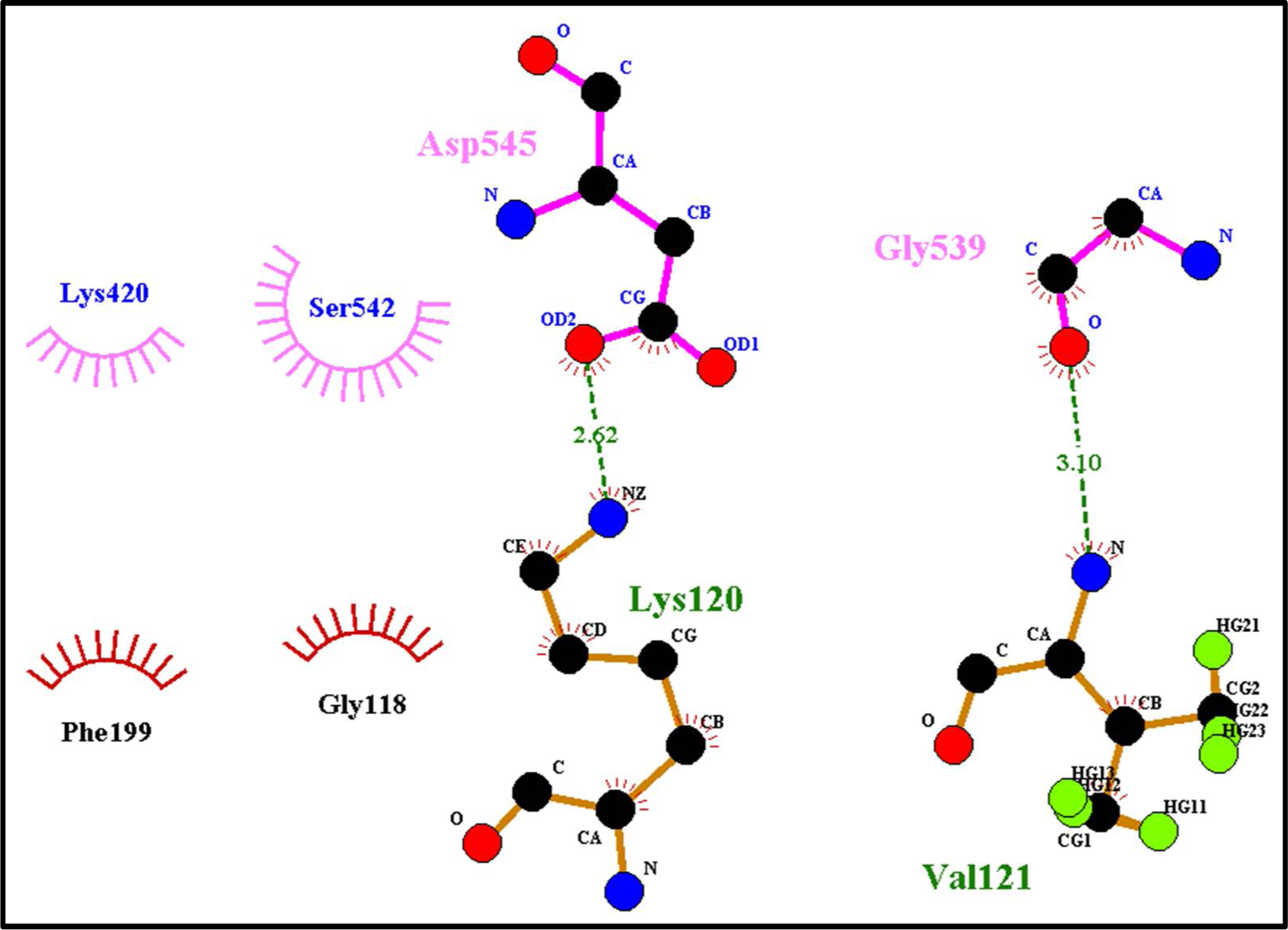
(**A**) Residue interaction network (RIN) showing the interactions between key residues of PFA0660w-PfHsp70-x docked complex. Nodes in blue and orange represent residues of PFA0660w and PfHsp70-x respectively. Different pairs of interacting residues (left to right in RIN plot) are mentioned. His114 of PFA0660w form maximum number of interaction network with Asp216, Lys217, Phe247, Ser416, Ala418 and Lys543 of PfHsp70-x. (**B**) Dimplot representing crucial residues forming interface of PFA0660w-PfHsp70-x docked complex. Green lines indicate the hydrogen bonds.

### Substrate binding domain

Atomic resolution structures of C-terminal peptide-binding fragments of type II Hsp40s have been solved for yeast sis1 (PDB ID: 1C3G), human Hdj1 (PDB ID: 2QLD) and *C. parvum* cgd2_1800 (PDB ID: 2Q2G) [34, 55, 61]. Yeast Sis1 and human Hdj1 are dimeric structures that carry two distinct domains that are comprised of β sheets. The hydrophobic depression forming peptide binding cavity resides in domain I [55, 60]. To get insight into the structural aspects of peptide binding clefts in case of *Plasmodium* species, we modeled the C-terminal peptide binding fragment of PFA0660w’s homologs in *Plasmodium*. The crystal structure of Hsp40 from *C. parvum* (PDB ID: 2Q2G) was used as a template by Swiss model tool. All model structures were refined and evaluated using different structure verification tools. The scores obtained suggest the reliability of these models for structure analysis (Table 5).

**Table 5:**
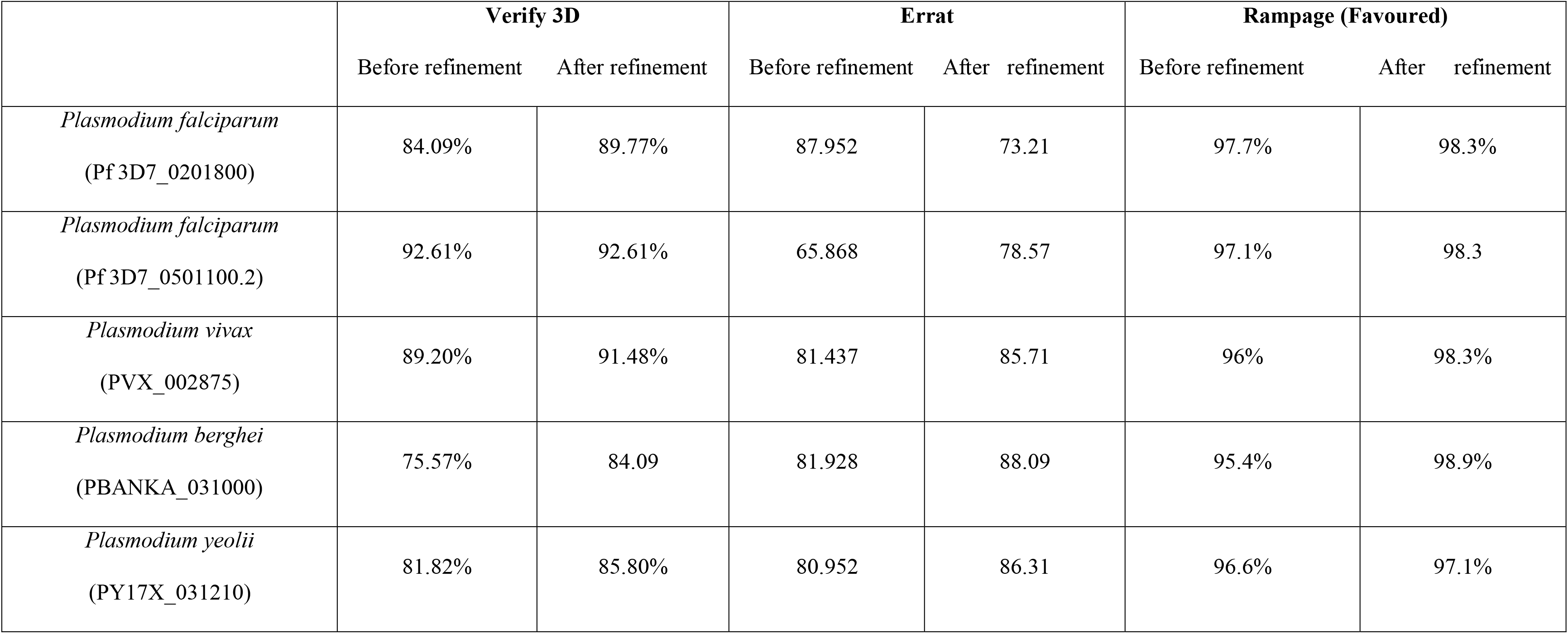
Validation scores for the models build for C terminal region of homologs of PFA0660w in *P. falciparum 3D7* and in other species.

Substrate binding pocket of PFA0660w and its homologs were computed using DoGSiteScorer server. Peptide binding cleft was identified by considering the residues (V184, L186, F201, I203 and F251) of yeast Sis1 reported to form substrate binding site and their corresponding residues in modeled and solved structures [55]. Analysis of hydrophobic depression forming substrate binding cleft in PFA0660w and its homologs suggests that they have bigger cavities as compared to yeast, human and *C. parvum* (Fig. 6; Table 6). Surface area and volume of cavities in PFA0660w and its homologs were more than that of yeast, human and *C. parvum* (Table 6). Area and volume of the cavity in PFA0660w was 314.81 Å^2^ and 199.19 Å^3^ respectively which was almost twice when compared with cavity size of *C. parvum* Hsp40. Peptide binding cleft in case of *P. vivax*, *P. berghe*i and *P. yeolii* was observed to be even larger than PFA0660w and its homologs in Pf3D7. Volume of cavity in *P. vivax* (PVX_002875), *P. berghei* (PBANKA_031000) and *P. yeolii* (PY17X_031210) were 350.14 Å3, 443.90 Å3, 496.90 Å3 respectively whereas in case of PFA0660w and its homologs in Pf3D7 (PF3D7_0201800, PF3D7_0501100.2), it was observed to be 199.19 Å3, 216.70 Å.3, 210.20 Å3 respectively. Multiple sequence alignment shows that hydrophobicity of amino acids involved in forming the peptide binding site of yeast Sis1 (V184, L186, F201, I203 and F251) is well conserved among *Plasmodium* strains (Fig. 7A). These residues are also conserved among type II Hsp40 family members (Fig. 7B) except for histidine which is present in place of valine in mouse, human and drosophila. However, V184 of yeast is conserved in Pf, *C. parvum and T. gondii*.

**Fig. 6.**
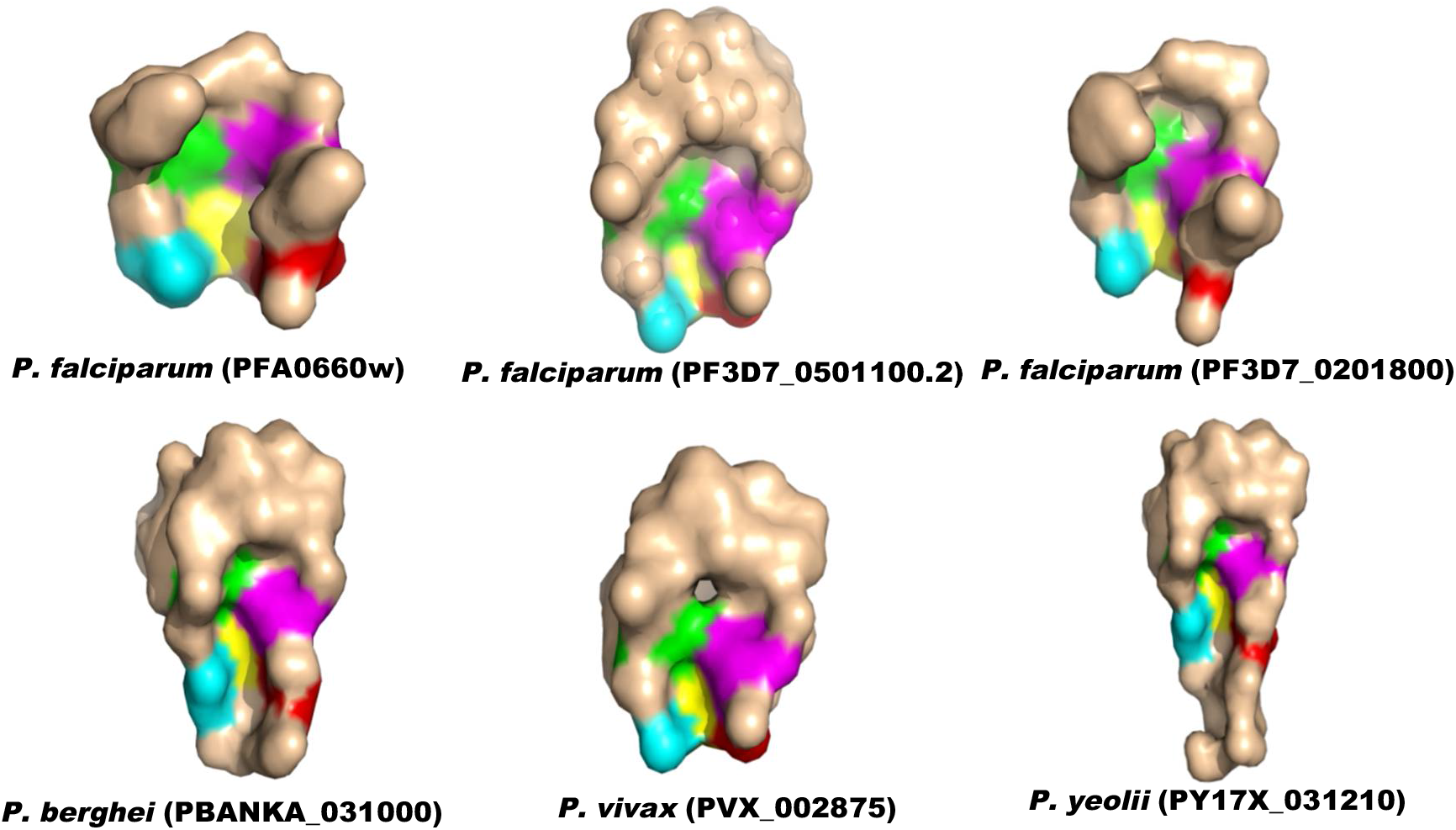
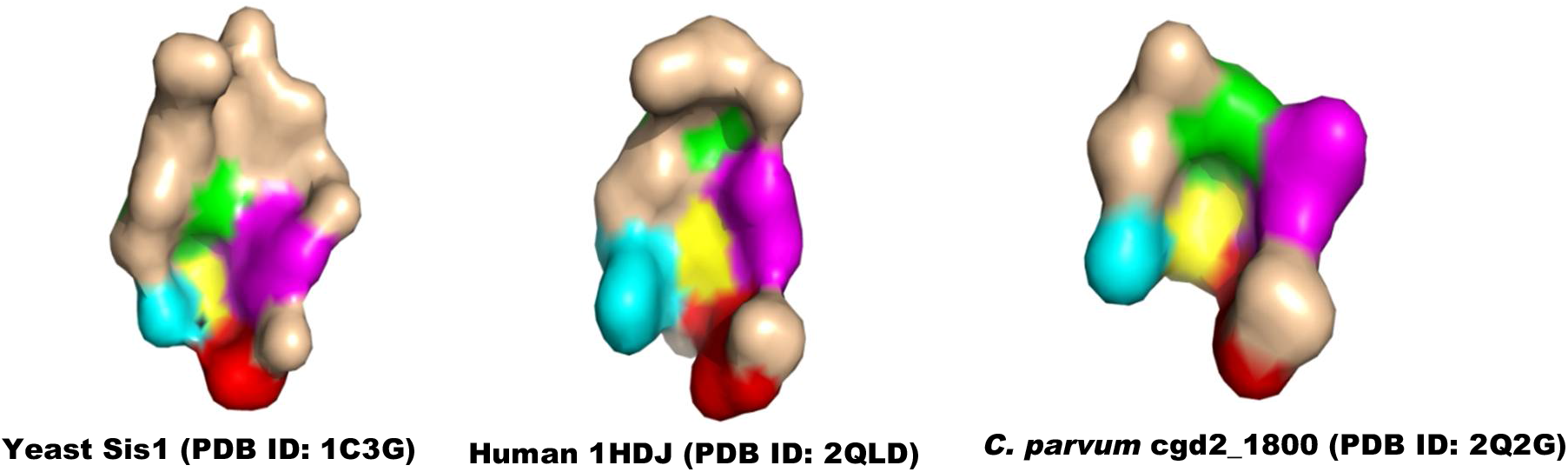
Surface representation of peptide binding clefts of modeled PFA0660w, its homologs in *Plasmodium* and crystal structures of yeast, human and *C. parvum* Hsp40s. Amino acid residues (V184, L186, F201, I203 and F251) known to form the peptide binding cleft of yeast Hsp40 and their corresponding residues are marked with cyan, green, magenta, red and yellow respectively.

**Fig. 7.**
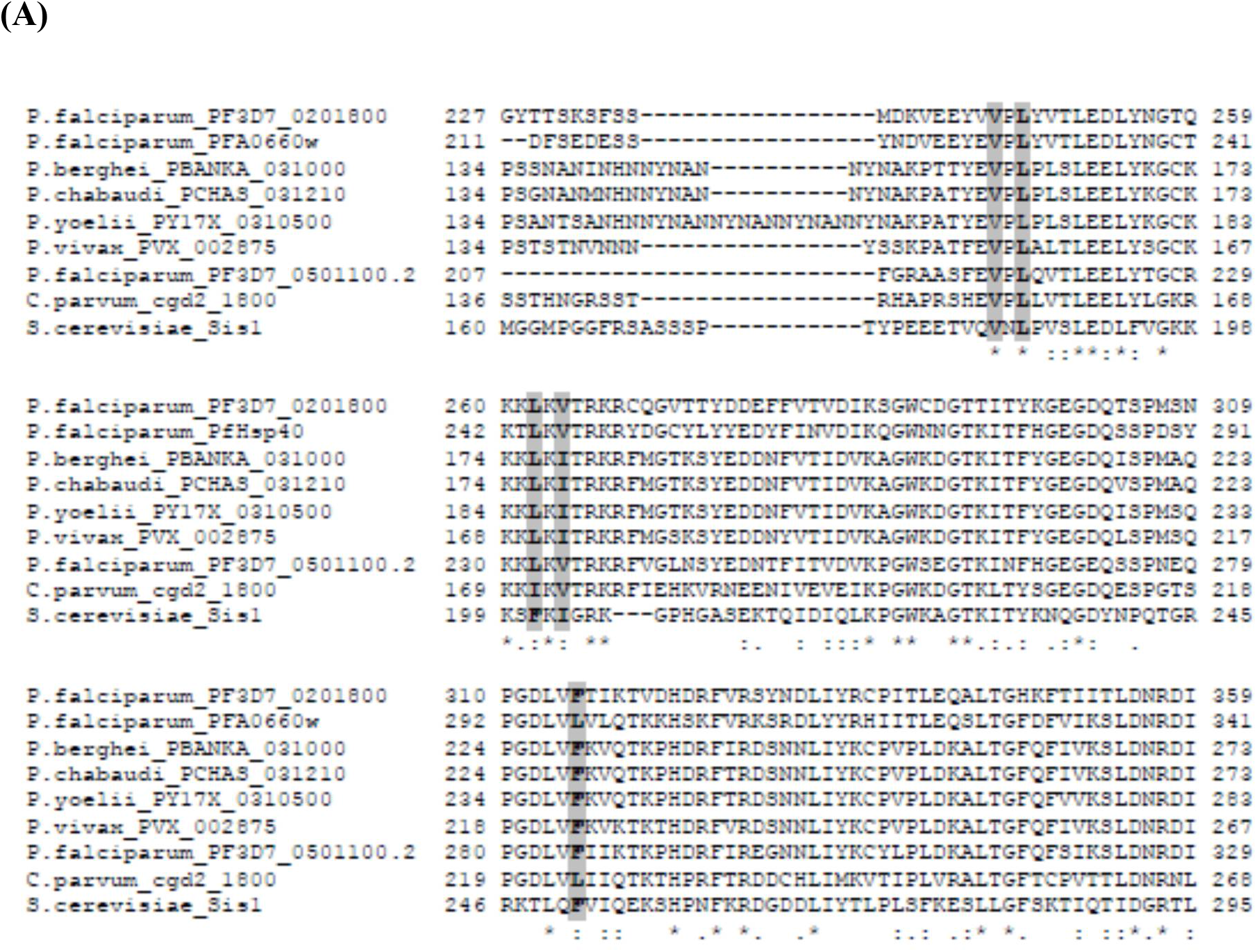
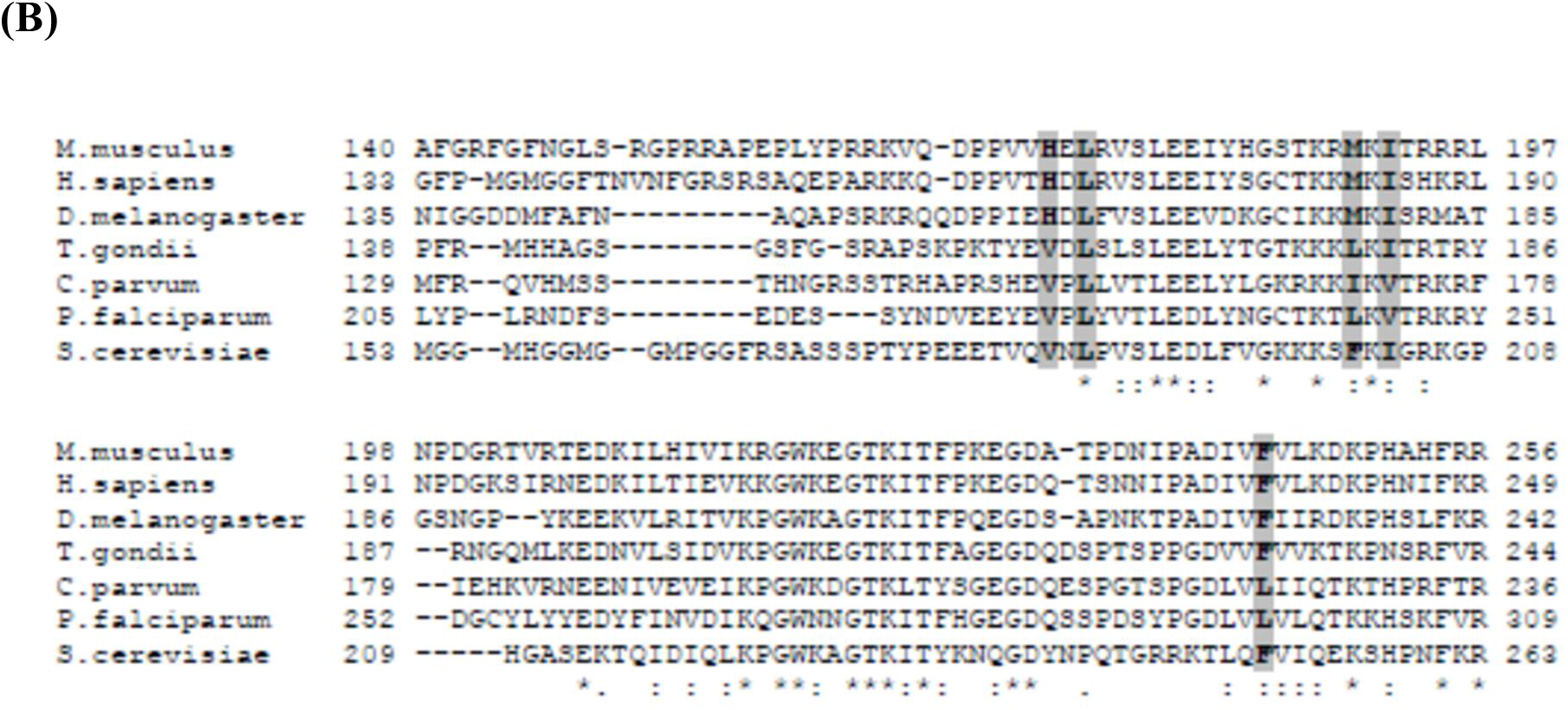
(**A**) Multiple Sequence alignment of type II Hsp40 homologs of PFA0660w in *Plasmodium falciparum 3D7* (PF3D7_0201800, PF3D7_0501100.2) and among different strains of *Plasmodium* viz. *P. vivax* (PVX_002875), *P. berghei* (PBANKA_0310000), *P. chabaudi* (PCHAS_031210) and *P. yeolii* (PY17X_0310500) (**B**) Multiple Sequence alignment of type II Hsp40 family members from *M. Musculus* (Hsp40-3), *H. sapiens* (Hdj1), *D. Melanogaster* (Droj-1), *T. gondii* (TgSis1), *S. Cerevisiae* (Sis1) and *C. parvum* (Cgd2_1800). The residues known to form substrate binding cavity in yeast Sis1 and their corresponding residues in other family members are marked with grey bars (A, B).

**Table 6:** Properties of peptide binding cleft of PFA0660w, its homologs in *Plasmodium falciparum 3D7* (PF3D7_0201800, PF3D7_0501100.2) and in different strains of *Plasmodium* including *P. vivax* (PVX_002875), *P. berghei* (PBANKA_0310000), *P. chabaudi* (PCHAS_031210), *P. yeolii* (PY17X_0310500), showing the comparison with solved structure of Hsp40 of yeast, human and *C. parvum*.

| Properties of pocket | *P. falciparum* 3D7 (PFA0660w) | *P. falciparum* 3D7 (PF3D7_0201800) | *P. falciparum* 3D7 (PF3D7_0501100.2) | *P. vivax* (PVX_002875) | *P. berghei* (PBANKA_031000) | *P. yeolii* (PY17X_031210) | Yeast (PDB ID:1C3G) | Human (PDB ID:2QLD) | *C. parvum* (PDB ID:2Q2G) |
| --- | --- | --- | --- | --- | --- | --- | --- | --- | --- |
| Volume [Å<sup>3</sup>] | 199.19 | 216.70 | 210.20 | 350.14 | 443.90 | 496.90 | 153.15 | 118.40 | 102.71 |
| Surface [Å<sup>2</sup>] | 314.81 | 421.27 | 377.45 | 582.66 | 728.56 | 835.64 | 301.68 | 286.26 | 139.57 |
| Depth [Å] | 9.91 | 12.06 | 14.57 | 14.87 | 18.36 | 23.99 | 10.54 | 9.62 | 7.16 |
| Enclosure | 0.22 | 0.21 | 0.18 | 0.17 | 0.15 | 0.17 | 0.30 | 0.24 | 0.32 |

### Effect of PfHsp70-x binding on substrate binding cavity of PFA0660w

To investigate whether binding of PFA0660w with PfHsp70-x influence its substrate binding domain, we interrogated the size of peptide binding cleft in representative docked structure obtained after MD simulations. Interestingly, the size of substrate binding cleft of PFA0660w in bound state was observed to be smaller as compared to unbound state. Area and volume occupied by the cavity of PFA0660w in PfHsp70-x bound state was 277.39 Å^2^ and 141.99 Å^3^ respectively whereas in unbound state it was 314.81 Å^2^ and 199.19 Å^3^ respectively. These results signify that binding of PFA0660w with PfHsp70-x may regulate the size of substrate binding cleft.

## Discussion

The number of Hsp40 proteins in Pf makes its chaperone machinery quite large and diverse. Inability to generate PFA0660w knock out parasite indicate that it may be essential for parasite survival in human host, making it an attractive candidate for anti-malarial drug discovery [7]. PFA0660w is a co-chaperone of PfHsp70-x and these co-localise to intracellular mobile structures termed as J-dots [29, 30]. These structures are implicated in the trafficking of major virulence factor ‘PfEMP1’ across the host cell [30] which further underscores their importance in malaria biology and makes them excellent targets for drug development. Considering vital role of Hsps during febrile episodes and in other processes like host cell remodelling, detailed structural analysis of PFA0660w-PfHsp70-x chaperone pair was conducted.

Structures of J domain and C-terminal peptide binding fragment of several Hsp40s have been solved experimentally [34, 55, 60]. However, structure for complete region including J domain, G/F and C-terminal domain has been determined only for *T. thermophilus* DnaJ (4J80), a typical prokaryotic type II Hsp40 [61]. To date, there is no experimentally solved structure for plasmodial chaperone. In the present study, an attempt was made to predict the model structure of complete conserved region of PFA0660w including J domain, G/F domain and SBD. As per our knowledge, the study provided for the first time a reliable model for the complete conserved region of any *Plasmodium* Hsp40 using the approach of advanced modeling. Multiple templates based 3D modeling has been reported to accurately align a single protein sequence simultaneously to multiple templates and build protein models with better quality than single-template based models [62].

Several reports demonstrated the mechanism of Hsp40-Hsp70 interaction using three dimensional structures of bacterial DnaJ-DnaK partners [18, 59]. Structure of DnaJ-DnaK complex using NMR reveals intermolecular interface to comprise of helix II of J domain and stretch of amino acids ranging from 206-221 of ATPase domain of DnaK [59]. Jiang *et al*. studied their interactions by covalently linking mammalian Hsc70 with auxilin; a neuronal specific Hsp40 [58]. Here, we have attempted to explore crucial residues forming the binding interface of PFA0660w-PfHsp70-x complex using their modeled structures. The model was obtained by applying protein–protein docking and molecular dynamics simulations followed by post MD interfacial analysis of the complex using RIN and Dimplot. MD simulation of docked complex revealed two distinct interaction sites in PFA0660w involving the residues of J domain and G/F region. Bipartite nature of binding of PFA0660w is consistent with previously known biochemical data that provide evidence for two distinct binding sites of DnaJ for DnaK [61, 63]. J domain residues of PFA0660w shows binding with residues located on ATPase domain and SBD of PfHsp70-x. Similar interface has been previously reported by Suh *et al*. where J domain of DnaJ forms interaction with ATPase domain and SBD of DnaK [56]. Experimental validation using NMR or crystallization studies of PFA0660w-PfHsp70 complex would give better insight into their interaction mechanism.

The C-terminal of Hsp40 proteins harbors the SBD for capturing the client protein and is further involved in dimerization [55, 60]. Hsp40s act as substrate scanning factors and selectors for Hsp70 which suggest that SBD play vital role in exhibiting their co-chaperone functions [19, 64]. Furthermore, it is known that Hsp40s show high affinity for substrates having aromatic and hydrophobic residues [64]. With the aim of getting comprehensive knowledge regarding the characteristic features of SBD, we modeled the C-terminus peptide binding fragments of PFA0660w’s homologs in *Plasmodium* species. All models obtained were refined and fully evaluated to show high quality structures that can be used for structural analysis. Interestingly, substrate binding cleft of PFA0660w and its homologs consisting of hydrophobic residues depict much bigger cavities as compared to their counterparts in yeast, human and *C. parvum*, suggesting their efficiency to capture bigger peptides. *In vitro* studies suggest that PFA0660w has a monomeric structure in solution unlike higher eukaryotes that are reported to exist as dimers [29]. The existence of bigger substrate binding cavities in *Plasmodium* species may support the monomeric structure of PFA0660w for holding substrate polypeptide while interacting and delivering it to PfHsp70-x. From our studies and previous data, we hypothesise a model representing PFA0660w’s interaction with substrate and PfHsp70-x. Our model suggests the independent ability of monomeric PFA0660w to capture peptide substrates owing to the presence of large substrate binding cleft (Fig. 8). On the contrary, Hsp40s in yeast, human and *C. parvum* having smaller peptide binding clefts dimerize to hold the polypeptide while exhibiting their chaperone function (Fig. 8). Large hydrophobic pockets of PFA0660w and its homologs may also provide specificity for peptide substrates. These distinguished structural features of SBD in PFA0660w and its homologs extend our knowledge in understanding the basis for holding the substrate peptides in *Plasmodium* species while performing their chaperone activity.

**Fig. 8.**
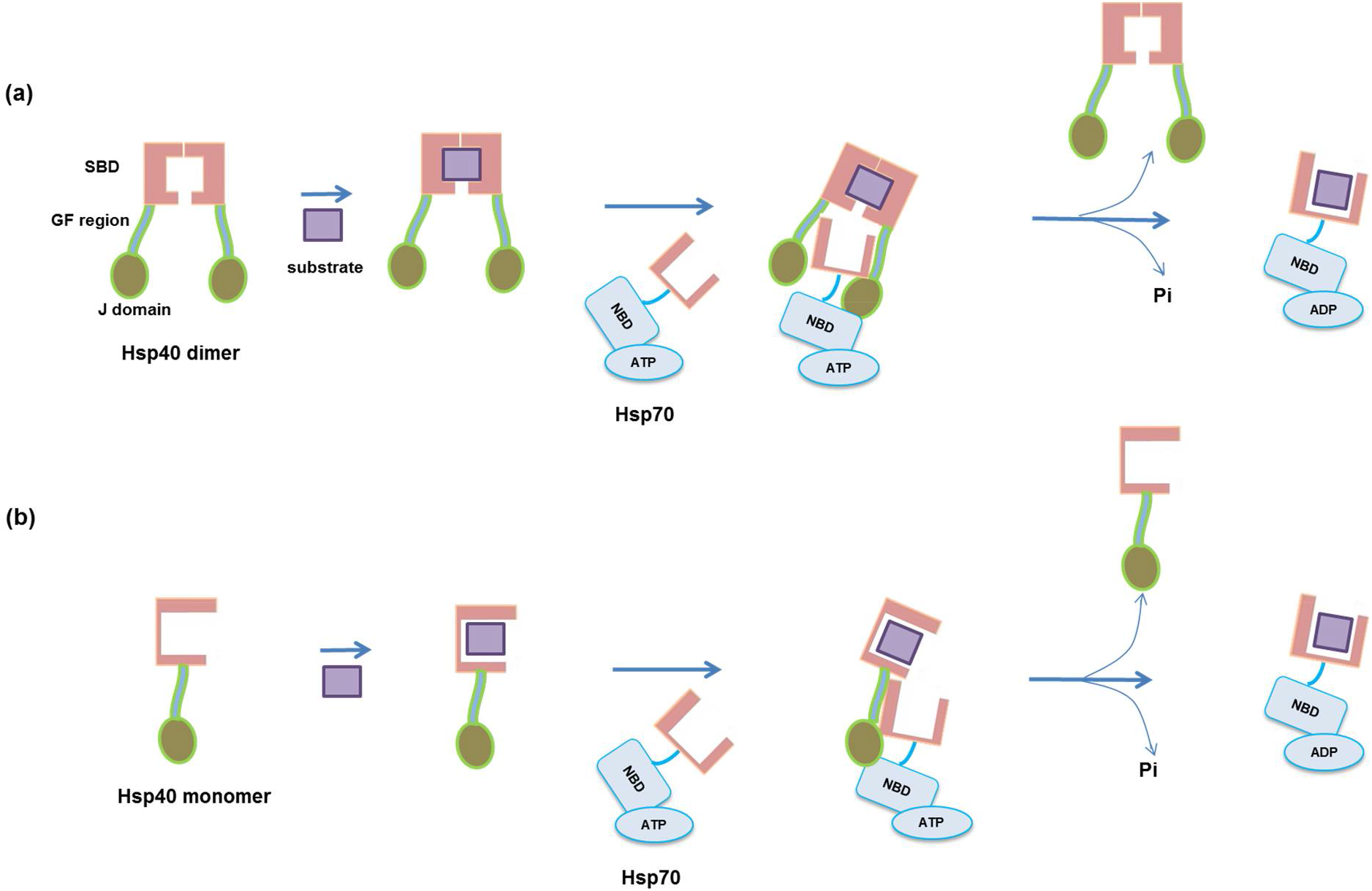
A hypothetical model for PFA0660w-PfHsp70-x interaction and a mechanism of substrate binding in PFA0660w. (**a**) Hsp40s in yeast, human and *C. parvum* have smaller peptide binding clefts and therefore form dimeric structures for holding the substrate while delivering it to their partner Hsp70. (**b**) PFA0660w carry bigger peptide binding pockets which may facilitate its monomeric form to hold the substrate independently while performing chaperone activity.

Representative structure of PFA0660w-PfHsp70-x complex obtained after MD simulation was also evaluated for computing the peptide binding cleft of PFA0660w in Hsp70 bound state. Interestingly, size of substrate binding pocket was found to reduce on binding with PfHsp70-x. These observations indicate that PFA0660w’s SBD exhibit significant flexibility to regulate the size of peptide binding cleft while interacting with PfHsp70-x.

Overall, analysis conducted in this study would contribute new knowledge regarding the structures and binding mechanism of an exported malarial chaperone pair ‘PFA0660w-PfHsp70-x’. Since PFA0660w is likely to be essential for parasite survival, structural characterization of this protein and interaction model with PfHsp70-x can be used as potential drug targets for the development of antimalarials.

## Conclusions

In the present study, we proposed for the first time a reliable three dimensional structure of entire conserved region of PFA0660w that includes J domain, G/F region followed by SBD. Docking and MD simulation studies of PFA0660w with its chaperone partner ‘PfHsp70-x’ provide insight into their interaction mechanism. Our results show that PFA0660w binds with PfHsp70-x in a bipartite manner. *In silico* studies reveal that substrate binding cavities in PFA660w and its homologs are bigger as compared to their counterparts in yeast, human and *C. parvum*. The existence of larger cavities in *Plasmodium* species provide support to our proposed model where monomeric PFA0660w holds polypeptide on its own while interacting and delivering it to PfHsp70-x. We also show that binding of PFA0660w with PfHsp70-x regulates the size of its substrate binding cavity. Knowledge gain in context of binding sites of such crucial proteins (PFA0660w) with their binding partner (PfHsp70-x) can lead to the development of new chemotherapeutics against malaria.

## Conflict of interest

Authors declare that they have no conflict of interest.

**Supplementary movie 1**: Trajectory movie of PFA0660w-PfHsp70-x docked complex for 2 ns demonstrating two distinct binding sites of PFA0660w for PfHsp70-x. Blue and red correspond to PFA0660w and PfHsp70-x respectively.

## Acknowledgements

PCM lab was supported by grant from Department of Biotechnology (DBT, Government of India). AB was a DBT-Senior Research Fellow (Department of Biotechnology-Govt. of India).

## Abbreviations

Pf: : *Plasmodium falciparum*
Hsp40: : Heat shock protein 40
Hsp70: : Heat shock protein 70
NBD: : nucleotide binding domain
SBD: : substrate binding domain
G/F: : Glycine-Phenylalanine

